# Cellular fitness is determined by ribosomal protein S12-mediated release of a truncated Xrp1

**DOI:** 10.1101/2025.10.29.685279

**Authors:** Bungo Kakemura, Hiroshi Kanda, Ryo Matsumoto, Shiori Ueda, Satoshi Yasuhara, Rina Nagata, Kiichiro Taniguchi, Shu Kondo, Keita Miyoshi, Tomoe Kobayashi, Kazuhiro Takeuchi, Kuniaki Saito, Makoto Matsuyama, Yasuhiro Murakawa, Tatsushi Igaki

## Abstract

Multicellular tissues require continuous optimization to maintain their structure and function by actively eliminating viable, yet unfit cells via a mechanism known as cell competition. During this process, unfit ‘loser’ cells commonly upregulate the C/EBP family transcription factor Xrp1, which causes their elimination, indicating that intracellular Xrp1 levels determine cellular fitness. However, despite its central role, the mechanism by which Xrp1 is upregulated in unfit cells remains unknown. Here, we show that Xrp1 is upregulated through a previously unrecognized post-transcriptional regulatory mechanism mediated by the ribosomal protein S12 (RpS12). Surprisingly, Xrp1 mRNA is abundantly expressed even in wild-type cells, yet it is not translated. This is due to an upstream open reading frame (uORF) located in the 5’UTR of the initial coding exon of *Xrp1* mRNA, which inhibits the translation of the main Xrp1 ORF. Intriguingly, in unfit cells, RpS12 causes splicing-mediated skipping of this initial exon, leading to the use of an alternate start codon that generates a short isoform of Xrp1 protein, causing cell death. Notably, protein structural analysis reveals that RpS12 is highly homologous to that of a spliceosomal component SNU13, suggesting the role of RpS12 in directly regulating alternative splicing of *Xrp1* mRNA. Our findings thus provide not only a mechanistic insights into how cellular fitness is determined but also how tissue optimization is achieved.

## Introduction

Multicellular communities have an intrinsic mechanism to continuously optimize its own structure and function by actively eliminating deleterious or unwanted cells via cell-cell communication. One such mechanism is cell competition, whereby viable but unfit cells (losers) are selectively eliminated from growing tissues when surrounded by fitter, wild-type cells (winners). Cell competition was first discovered in *Drosophila* (Morata and Ripoll, 1975) and was later shown to occur in vertebrates (Hogan *et al*., 2009; Clavería *et al*., 2013; Sancho *et al*., 2013; Akieda *et al*., 2019; Hashimoto and Sasaki, 2019), suggesting an evolutionarily conserved mechanism of multicellular optimization. Studies in different model systems have suggested that cell competition plays important roles in tumor elimination, cancer expansion, selection of fitter stem cells, and animal aging (reviewed in (Clavería and Torres, 2016; Di Gregorio, Bowling and Rodriguez, 2016; Merino, Levayer and Moreno, 2016; Baker, 2020; Vishwakarma and Piddini, 2020; Nagata and Igaki, 2024)).

Studies in *Drosophila* imaginal epithelia have revealed that cell competition is triggered by a variety of gene mutations such as in a *ribosomal protein* (Rp) gene (also known as ‘*Minute*’ mutation) (Morata and Ripoll, 1975), an E3 ligase *mahjong* (*mahj*) (Tamori *et al*., 2010) a DEAD box RNA helicase *Helicase at 25E* (*Hel25E*) (Nagata *et al*., 2019), ER proteins such as *wollknaeuel* (*wol*) or *Calreticulin* (*Calr*)(Ochi *et al*., 2021) translation factors such as *eIF4G*, *eIF5A*, *eIF6*, *eEF1a*, and *eEF2* (Kiparaki *et al*., 2022), and apicobasal polarity genes such as *scribble* (*scrib*) or *discs large* (*dlg*) (Brumby and Richardson, 2003; Igaki *et al*., 2009) when mutant cells are surrounded by wild-type cells. In addition, differences in intracellular Myc levels between cells can trigger cell competition, as cells with higher Myc expression eliminate neighboring cells with lower Myc expression by inducing cell death (de la Cova *et al*., 2004; Moreno and Basler, 2004). These mutations, apart from apicobasal polarity mutations and relative Myc expressions, cause cell competition in a C/EBP family transcription factor Xrp1-dependent manner (Baillon *et al*., 2018; Lee *et al*., 2018). These loser mutants commonly upregulate Xrp1, which is sufficient to induce cell death via upregulation of proapoptotic genes such as *reaper* (*rpr*) and *head involution defective* (*hid*) (Baillon *et al*., 2018). Thus, the intracellular level of Xrp1 protein determines ‘cellular fitness’, as elevation of Xrp1 makes cells unfit and thereby causing cell competition with neighboring wild-type cells. Xrp1 can regulate its own transcription via a positive autoregulatory loop (Baillon *et al*., 2018), through heterodimerization with a C/EBP transcription factor Irbp18 (Blanco, Cooper and Baker, 2020). Both Xrp1 and Irbp18 are upregulated in loser cells and are required for the induction of Xrp1. Despite extensive studies, however, the mechanism by which Xrp1 protein is upregulated in loser cells still remains unknown.

Genetic studies have shown that the phosphorylation of the alpha subunit of eukaryotic translation initiation factor 2 (eIF2α) appears to be a key event that leads to the upregulation of Xrp1 (Kiparaki *et al*., 2022). The role of eIF2α has been previously described as the integrated stress response that suppresses global protein synthesis while selectively allowing the translation of specific genes, such as activating transcription factor 4 (ATF4) (Costa-Mattioli and Walter, 2020). Interestingly, elevated levels of phosphorylated eIF2α (p-eIF2α) are sufficient to induce the transcriptional upregulation of Xrp1 (Ochi *et al*., 2021; Kiparaki *et al*., 2022). Moreover, loser cells commonly exhibit increased p-eIF2α levels through Xrp1 activity (Langton *et al*., 2021; Ochi *et al*., 2021; Kiparaki *et al*., 2022), suggesting the existence of a positive feedback loop between p-eIF2α and Xrp1 that promotes cell death. Intriguingly, although overexpression of ATF4 is sufficient to induce Xrp1 transcription, ATF4 is not required for Xrp1 upregulation in *Minute* loser cells (Langton *et al*., 2021), suggesting that p-eIF2α promotes Xrp1 transcription in these cells through an unknown, ATF4-independent mechanism.

A genetic screen to determine potential upstream molecule required for the induction of Xrp1 identified the ribosomal protein S12 (RpS12)(Kale *et al*., 2018). Intriguingly, missense (G97D) mutation of *RpS12*, but not null mutations, suppressed Xrp1 induction and subsequent cell death in *Minute* cell competition (Kale *et al*., 2018), suggesting that RpS12^G97D^ does not act as a loss-of-function mutation. Nevertheless, exactly how the *RpS12^G97D^* mutation suppresses Xrp1 induction and how RpS12 precisely contributes to Xrp1 induction still remains unknown.

## Results

### RpS12 upregulates Xrp1 at the post-transcriptional level

In *Drosophila* imaginal discs, many unfit cells (though not all) are eliminated through the upregulation of Xrp1 when surrounded by wild-type cells (Figure S1A and B). As this process can be blocked by the knockdown of *Xrp1* (Figure S1C-H), this indicates that the upregulation of Xrp1 plays a central role in their elimination. To understand the mechanism by which Xrp1 is upregulated in unfit cells, we first investigated the role of RpS12 as a potential upstream regulator of Xrp1 induction in loser cells (Kale *et al*., 2018; Lee *et al*., 2018; Ji *et al*., 2019). We found that overexpression of RpS12 in *Drosophila* larval wing disc was sufficient to upregulate both *Xrp1* transcription and Xrp1 protein levels (Figure 1A and B, quantified in 1D and E). Similarly, the overexpression of ATF4, which has been previously reported to upregulate *Xrp1* transcription (Langton *et al*., 2021), indeed also led to the upregulation of transcription, but also protein levels of Xrp1 (Figure 1C, quantified in 1D and E). Considering that Xrp1 transcription can be induced by its own autoregulation (Baillon *et al*., 2018; Blanco, Cooper and Baker, 2020),we next performed the same analysis using *Xrp1* mutant animals, to prevent the production of Xrp1 proteins and disrupts its autoregulation. Indeed, overexpression of RpS12 or ATF4 did not upregulate Xrp1 protein levels in *Xrp1-lacZ/Xrp1^m2-73^* mutant wing discs (Figure 1F-H, specifically F’’, G’’, and H’’, quantified in 1I and J). In contrast, overexpression of ATF4 in these mutant tissues elevated the transcriptional reporter *Xrp1-lacZ* (Figure. 1H’, quantified in 1I), indicating that ATF4 transcriptionally upregulates Xrp1. Intriguingly, RpS12 overexpression did not upregulate the *Xrp1* transcription activity (Figure. 1G’, quantified in 1I), which suggests that RpS12 induces Xrp1 likely at the post-transcriptional level.

**Fig. 1.**
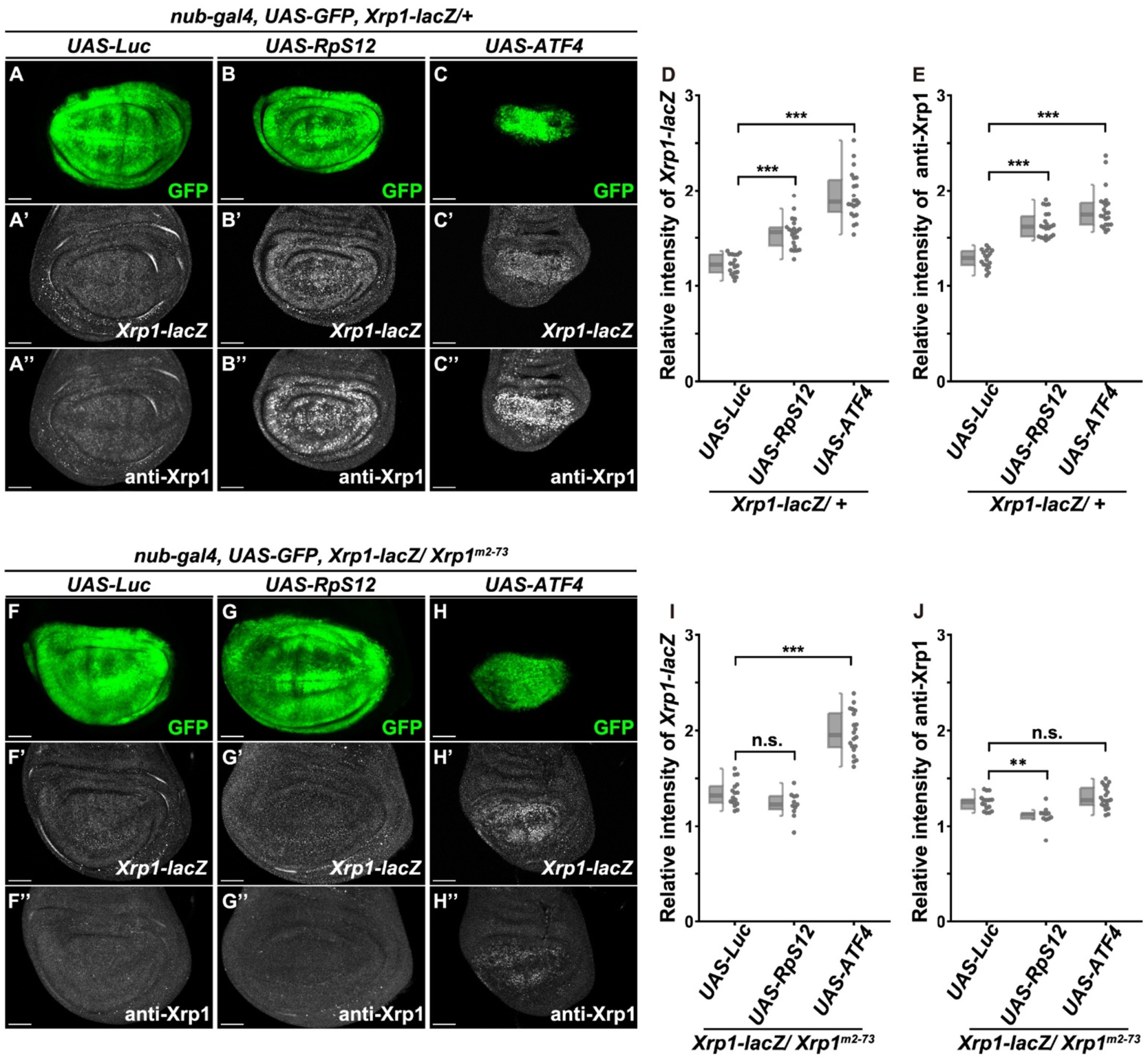
RpS12 upregulates Xrp1 at the post-transcriptional level. (A-C) Wing discs of *Xrp1-lacZ/+* background larvae, expressing Luc (control), RpS12, or ATF4 under *nub-Gal4* control, stained with anti-Xrp1 and anti-β-gal antibodies. Scale bars, Scale bars, 50 μm. Confocal images were acquired with a confocal microscope ZEISS LSM 880 (Carl Zeiss) under the control of ZEN Blue (Carl Zeiss). (D) Boxplot overlaid with dot plot represents the relative intensity of *Xrp1-lacZ* in GFP positive cells normalized by the signal in GFP negative region in the pouch for Luc (n=18), RpS12 (n=22), ATF4 (n=22) samples. Each plot corresponds to the raw data. Statistical significance is shown as follows:***p < 0.001; Kruskal-Wallis test followed by the Steel-Dwass test. (E) Boxplot overlaid with dot plot represents the relative intensity of anti-Xrp1 in GFP positive cells normalized by the signal in GFP negative region in the pouch for Luc (n=18), RpS12 (n=22), ATF4 (n=22) samples. Each plot corresponds to the raw data. Statistical significance is shown as follows:***p < 0.001, n.s. (not significant) p>0.05; Kruskal-Wallis test followed by the Steel-Dwass test. (F-H) Wing discs of *Xrp1-lacZ/ Xrp1^m2-73^* background larvae, expressing Luc (control), RpS12, or ATF4 under *nub-Gal4* control, stained with anti-Xrp1 and anti-β-gal antibodies. Scale bars, 50 μm. Confocal images were acquired by a confocal microscope ZEISS LSM 880 (Carl Zeiss) under the control of ZEN Blue (Carl Zeiss). (I) Boxplot overlaid with dot plot represents the relative intensity of *Xrp1-lacZ* in GFP positive cells normalized by the signal in GFP negative region in the pouch for Luc (n=16), RpS12 (n=11), ATF4 (n=20) samples. Each plot corresponds to the raw data. Statistical significance is shown as follows: ***p < 0.001, n.s. (not significant) p>0.05; Kruskal-Wallis test followed by the Steel-Dwass test. (J) Boxplot overlaid with dot plot represents the relative intensity of anti-Xrp1 in GFP positive cells normalized by the signal in GFP negative region in the pouch for Luc (n=16), RpS12 (n=11), ATF4 (n=20) samples. Each plot corresponds to the raw data. Statistical significance is shown as follows: **p < 0.01, n.s. (not significant) p>0.05; Kruskal-Wallis test followed by the Steel-Dwass test.

### Loser cells upregulate Xrp1-S but not Xrp1-L protein levels

Based on our observations that RpS12 post-transcriptionally upregulates Xrp1, we next analyzed the Xrp1 protein isoforms expressed in loser cells. At least seven splice isoforms of *Xrp1* mRNA have been predicted to produce either long (L) or short (S) form of Xrp1 proteins (Baillon *et al*., 2018; Lee *et al*., 2018). Xrp1-L or Xrp1-S is translated from *Xrp1* mRNA isoform with or without exon 4, respectively. We performed a full-length RNA sequencing (RNA-seq) of Xrp1 in wild-type wing discs and found that *Xrp1* mRNAs indeed include seven splice isoforms (Figure 2A and S2). The RNA-seq data revealed that a long form of *Xrp1* mRNA bearing exon 4, *Xrp1-L*, is a major transcript from the Xrp1 gene (Figure S2). We then analyzed Xrp1 proteins expressed in wild-type or *Minute*/+ wing discs. Surprisingly, Xrp1-L protein was almost undetectable in both wild-type and unfit *Minute*/+ discs such as *RpS18^-/+^* or *RpS3^-/+^* discs (Figure 2B). Instead, the level of truncated Xrp1-S protein was strongly upregulated in *Minute*/+ discs (Figure 2B). This suggests that Xrp1-S protein, but not Xrp1-L protein, plays a crucial role in eliminating loser cells. Consistently, elimination of *RpL19^-/+^*clones by cell competition (Kiparaki *et al*., 2022) (Figure 2C) was significantly suppressed by introducing *Xrp1-short*-specific mutations (*Xrp1^08^/Xrp1^0^*^8^,referred to as *ΔXrp1-S*) (Baillon *et al*., 2018) in loser clones (Figure 2D, quantified in Figure 2I). Moreover, *Xrp1-long*-specific mutations, which was generated by deleting 5 bases in the coding region of exon 4 (*Xrp1^041^/Xrp1^041^,* indicated as *ΔXrp1-L,* see Methods) (Figure S3), showed significantly weaker suppression of cell competition compared to *ΔXrp1-S* (Figure 2E, quantified in Figure 2I). Similarly, *Hel25E*-induced cell competition was significantly suppressed in the *ΔXrp1-S* background while *ΔXrp1-L* only partially suppressed it (Figure 2G and H, quantified in Figure 2J). Overexpression of either Xrp1-L or Xrp1-S caused cell death in the imaginal disc (Figure S3). These data suggest that unfit loser cells upregulate Xrp1-S protein, as the major cause of loser’s cell death.

**Fig. 2.**
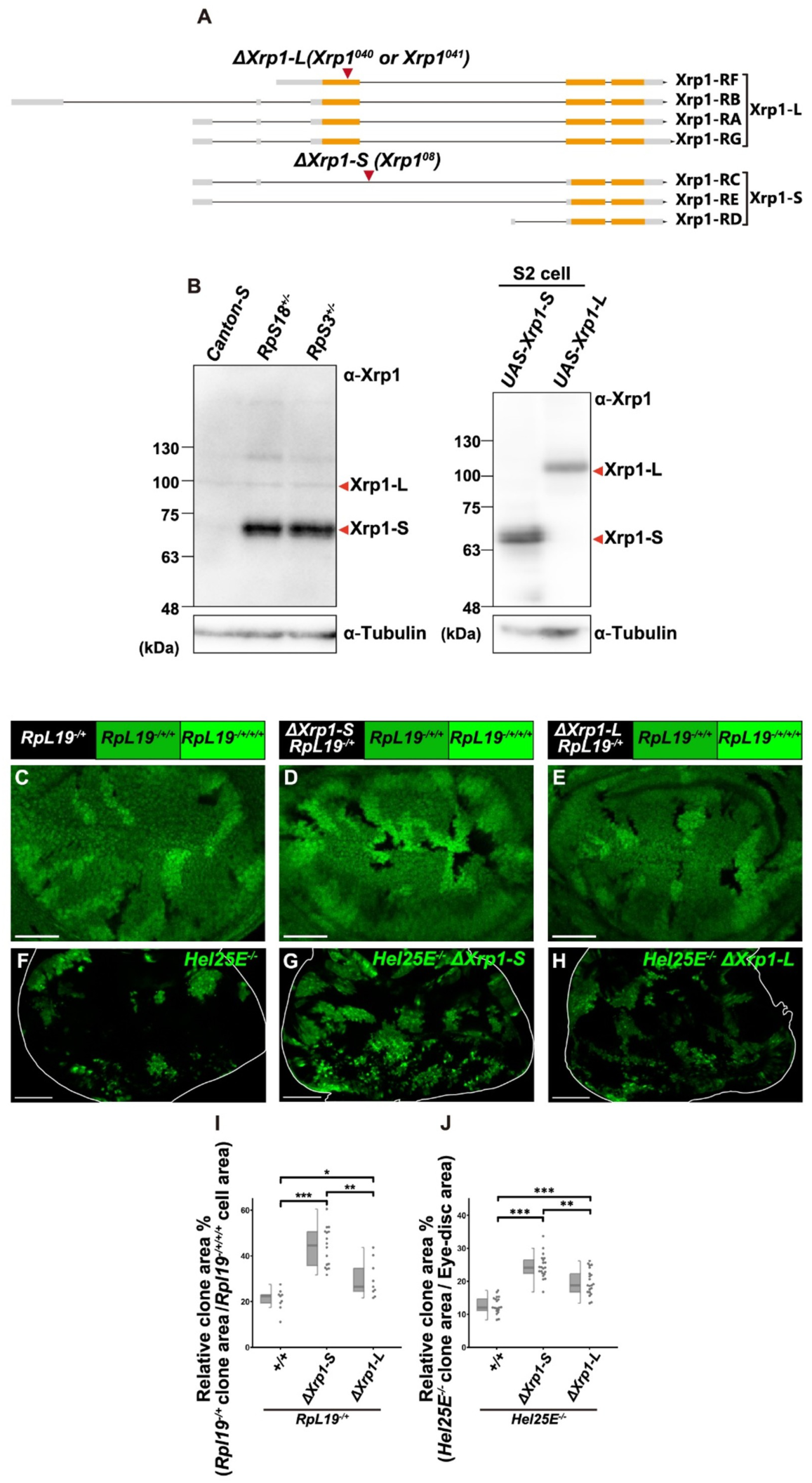
Loser cells upregulate Xrp1-S to drive cell competition. (A) Illustration of Xrp1 transcript isoforms. *Xrp1-RA, RB, RF,* and *RG* are *Xrp1 mRNA* long isoforms. *Xrp1-RC, RD,* and *RE* are *Xrp1 mRNA* short isoforms. (B) Wing discs of third instar larvae of each genotype or S2 cells transfected with each vector were subjected to Western blot analysis using anti-Xrp1 antibody or anti-Tubulin antibody. (C-E) Wing discs bearing *hs-FLP* induced *RpL19^-/+^*(C), *RpL19^-/+^+Xrp1^08/08^ (ΔXrp1-S)* (D) or *RpL19^-/+^+Xrp1^041/041^(ΔXrp1-L)* (E) cells. Scale bars, 50 μm. Confocal images were acquired with a confocal microscope ZEISS LSM 880 (Carl Zeiss) under the control of ZEN Blue (Carl Zeiss). (F-H) Eye discs bearing *ey-FLP* induced *Hel25E*^-/-^ clones in wild-type background (F), in *Xrp1^08/08^ (ΔXrp1-S)* background (G) or in *Xrp1^040^/ Xrp1^041^(ΔXrp1-L)* background (H). Scale bars, 50 μm. Confocal images were acquired with a Laser Scanning Confocal Microscope TCS SP8 on DMi8 inverted microscope (Leica Microsystems). (I) Box plot overlaid with dot plot represents the relative *RpL19^-/+^* clone area per *RpL19^-/+/+/+^* clone for *RpL19* (n=9), *RpL19Δ+Xrp1-S* (n=16), *RpL19*+*ΔXrp1-L* (n=9). Each plot corresponds to the raw data. Statistical significance is shown as follows: *p < 0.05, **p < 0.01, ***p < 0.001; Kruskal-Wallis test followed by the Steel-Dwass test. (J) Box plot overlaid with dot plot represents the relative GFP positive clone area per eye disc area for *Hel25E^-/-^* (n=19), *Hel25E^-/-^ ΔXrp1-S* (n=21), *Hel25E^-/-^ ΔXrp1-L* (n=21) cells. Each plot corresponds to the raw data. Statistical significance is shown as follows:**p < 0.01, ***p < 0.001; Kruskal-Wallis test followed by the Steel-Dwass test.

### Loser cells upregulate *Xrp1-S* mRNA variant

We next examined how Xrp1-S protein is upregulated in loser cells. Given that Xrp1 is post-transcriptionally upregulated by RpS12 (Figure 1), we hypothesized that Xrp1-S protein is induced by increased *Xrp1-S* mRNA isoform via RpS12, potentially by alternative splicing of *Xrp1* mRNA, in *Minute*/+ cells. To test this possibility, we performed a full-length RNA-seq of *Xrp1* mRNA in wild-type and *RpS18^-/+^* wing discs and found that a short variant *Xrp1-RC* is significantly elevated in *RpS18^-/+^* discs (Figure 3A). The RNA-seq also revealed that a long variant *Xrp1-RG* was the major variant of *Xrp1* transcript and that exon 4-skipping splicing that generates *Xrp1-RC* and *Xrp1-RE* (denoted as ‘splicing index’) was significantly increased in *RpS18^-/+^* cells (Figure 3B). We further confirmed this by performing semi-quantitative RT-PCR analysis and found that *Xrp1-RC* is significantly increased in *RpS18^-/+^* or *RpS3^-/+^* unfit cells compared to wild-type cells (Figure 3C and 3D, quantified in Figure 3E, Figure S4). This increase in *Xrp1-RC* levels in *RpS18^-/+^* cells was cancelled in *RpS12^G97D^*homozygous mutant background (Figure 3D, quantified in Figure 3E), indicating that RpS12 plays a crucial role in the induction of *Xrp1-S* variant in *Minute*/+ cells. This was further validated by using a different primer set that amplifies *Xrp1-RC* and *Xrp1-RE* short isoforms (Figure 3F, quantified in Figure 3G, Figure S4). We also analyzed a short variant *Xrp1-RD*, which is generated in a splicing-independent manner, and found that its level was unchanged in *RpS18^-/+^* cells (Figure S4). Western blot analysis revealed that the induction of Xrp1-S protein in *RpS18^-/+^* cells was abolished in *RpS12^G97D^* mutant background (Figure 3H). Consistent with these data, overexpression of RpS12 upregulated both *Xrp1-S* mRNA and Xrp1-S protein levels (Figure S4). These data suggest that loser cells specifically elevate *Xrp1-S* mRNA variant via RpS12, which produces a truncated Xrp1-S protein that causes cell death. Strikingly, we found by Foldseek analysis (van Kempen *et al*., 2023) that the whole protein structure of RpS12 is highly homologous to that of small nuclear ribonucleoprotein 13 (SNU13)/NHP2-like protein 1 (NHP2L1), a component of the U4/U6.U5 tri-snRNP complex of the spliceosome (Figure 3I). These data suggest that unfit cells elevate Xrp1 protein levels by selectively upregulating *Xrp1-S* mRNA via RpS12-mediated splicing of *Xrp1* mRNA.

**Fig. 3.**
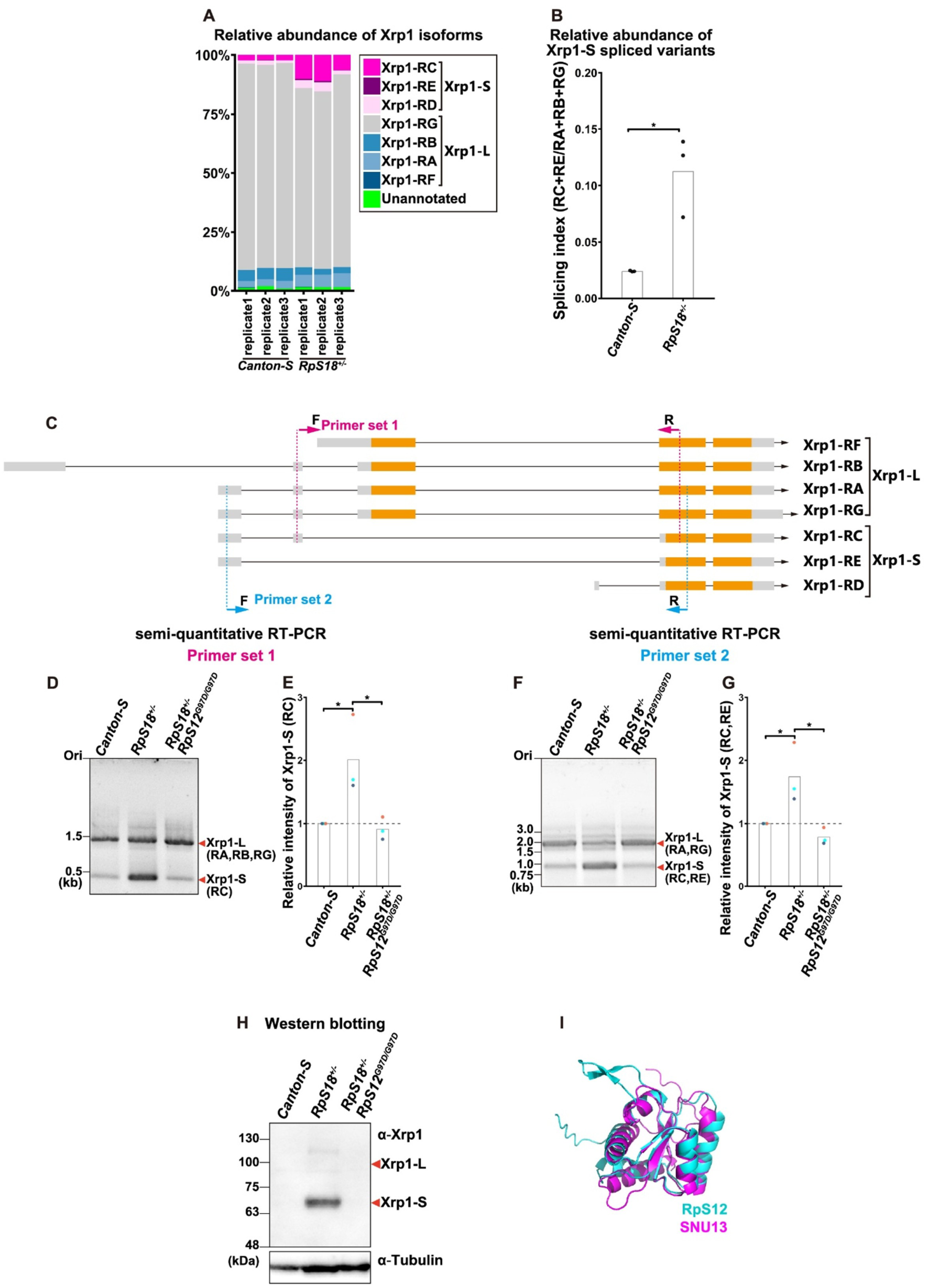
Loser cells upregulate *Xrp1-S* variant via RpS12. (A) Stacked bar plot showing the relative abundance of Xrp1 isoforms in individual biological replicates. The first three bars represent wild-type (WT) samples, and the last three bars represent S18 samples. Each colored segment indicates the percentage of a specific isoform relative to the total Xrp1 transcripts. (B) Bar plot overlaid with dot plot represents the relative intensity of the splicing index. The splicing index was calculated as the ratio of the *Xrp1-RC* and *Xrp1-RE* isoforms to the sum of the Xrp1-RA, –RB, and *-RG* isoforms. (*Xrp1-RC* + *Xrp1-RE* / [*Xrp1-RA* + *Xrp1-RB* + *Xrp1-RG*]). Each dot plot corresponds to the raw data. White bars show the average. Statistical significance is shown as follows: *p < 0.05; Welch’s t-test. (C) Illustration for primer-sets used for the amplification of *Xrp1* mRNA. (D) Semi-qRT-PCR products amplified using primer set 1 from cDNA derived from *Canton-S* (left), *RpS18^+/-^* (center), and *RpS18^+/-^* with *RpS12^G97D/G97D^*(right). (E) Bar plot overlaid with dot plot represents the relative intensity of *Xrp1-S* (RC) normalized by control sample. Each dot plot corresponds to the raw data, and each plot color corresponds to each replicate. White bars show the average. Statistical significance is shown as follows: *p < 0.05; one-way ANOVA followed by Tukey HSD test. (F) Semi-qRT-PCR products amplified using primer set 2 from cDNA derived from *Canton-S* (left), *RpS18^+/-^* (center), and *RpS18^+/-^* with *RpS12^G97D/G97D^*(right). (G) Bar plot overlaid with dot plot represents the relative intensity of *Xrp1-S* (RC, RE) normalized by control sample. Each dot plot corresponds to the raw data, and each plot color corresponds to each replicate. White bars show the average. Statistical significance is shown as follows: *p < 0.05; one-way ANOVA followed by Tukey HSD test. (H) Wing discs of third instar larvae of each genotype were subjected to Western blot analysis using anti-Xrp1 antibody or anti-Tubulin antibody. (L) Structure of *Drosophila* RpS12 aligned to *Human* SNU13/ NHP2L1

### Xrp1-L translation is normally inhibited by uORF, which is removed in loser cells

Our data indicate that, while *Xrp1-S* mRNA is upregulated in unfit *Minute*/+ cells, *Xrp1-L* mRNA is abundantly expressed even in wild-type cells (Figure. 3A). Notably, however, Xrp1-L protein was undetectable in these cells (Figure. 2B and 3H), suggesting that translation of *Xrp1-L* mRNA is compromised in some manner. One possible mechanism by which translation of a coding sequence can be specifically attenuated is the presence of an ectopic start codon upstream of the main ORF (Barbosa, Peixeiro and Romão, 2013). Indeed, we found an ATG codon with a strong Kozak sequence located at 208 nucleotides upstream of the *Xrp1-L* mRNA start codon in the *Xrp1-L-*specific exon 4 (Figure. 4A), as predicted previously (Brown *et al*., 2021). This uORF consists of 864 nucleotides that potentially produces 288 amino acid protein, which we named Xrp1-uORF 288 peptide (Xu288), thereby disrupting the translation of downstream *Xrp1-L* main ORF (Figure. 4B). These lines of evidence suggest that translation of *Xrp1-L* mRNA may normally be inhibited by the presence of uORF. To test this possibility, we used mScarlet as a fluorescent translation reporter in *Drosophila* S2 cells. As expected, the 5’-UTR of *Xrp1-L* mRNA, which includes the Xu288 start codon, inhibited translation of the downstream mScarlet ORF (Figure. 4C and D, quantified in Figure. 4G). This inhibition of mScarlet translation was abolished when the exon 4 5’-UTR or uORF start codon was removed from the reporter (Figure. 4C, E and F, quantified in Figure. 4G). These findings suggest that translation of *Xrp1-L* mRNA is normally blocked by the presence of Xu288 ORF.

**Fig. 4.**
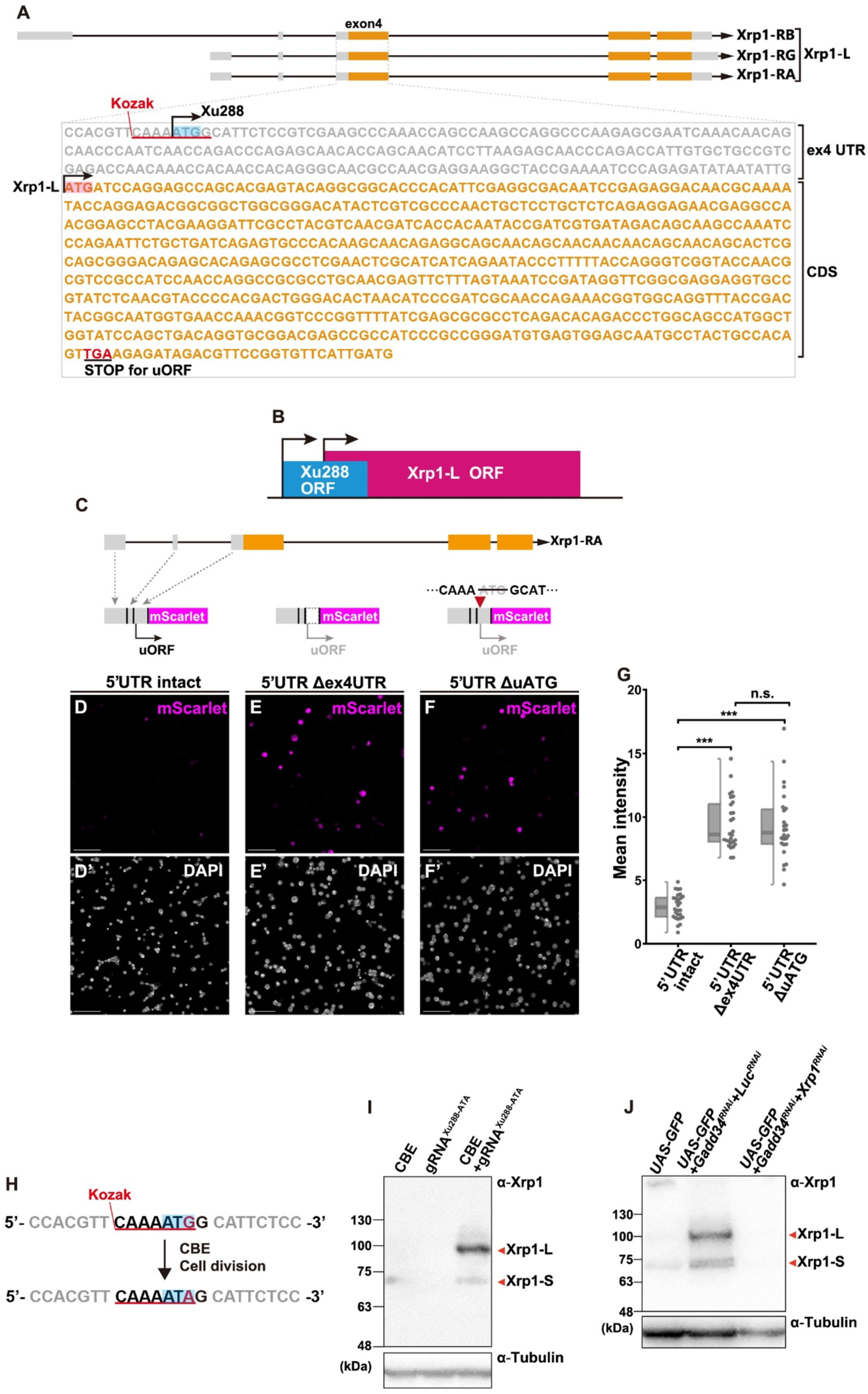
Xrp1-L translation is normally inhibited by uORF. (A) Schematic of Xrp1 reading frames. The uORF overlaps the main coding frame, and a strong Kozak motif lies immediately upstream of the Xrp1-L ORF. (B) Schematic diagram of Xu288 ORF and Xrp1-L ORF. (C) Illustration of pUAST-attB vector expressing mScarlet fused to 5’UTR of Xrp1-RA isoform. (D-F) S2 cells transfected with pUAST-mScarlet vector (D), pUAST-mScarlet vector lacking the 5’UTR region in exon 4 (E), pUAST-mScarlet vector lacking the ATG codon of Xu288 (F). Scale bars, 50 μm. Confocal images were acquired with Laser Scanning Confocal Microscope STELLARIS 5 WLL on DMi8 inverted microscope (Leica Microsystems). (G) Boxplot overlaid with dot plot represents the mean fluorescence intensity divided by cell area for intact (n=28), 5’UTRΔex4UTR (n=28), or ΔATG (n=26). Each plot corresponds to each image. Statistical significance is shown as follows: ***p < 0.001, n.s. p >0.05; Kruskal-Wallis test followed by the Steel-Dwass test. (H) Schematics of ATG-to-ATA editing in the uORF using CBE and gRNA^Xu288-ATA^. The Kozak motif is underlined; the target ATG codon is highlighted in blue. (I) S2 cells transfected with each vector were subjected to Western blot analysis using anti-Xrp1 antibody or anti-Tubulin antibody. (J) Wing discs of third instar larvae of each genotype were subjected to Western blot analysis using anti-Xrp1 antibody.

**Fig. 5.**
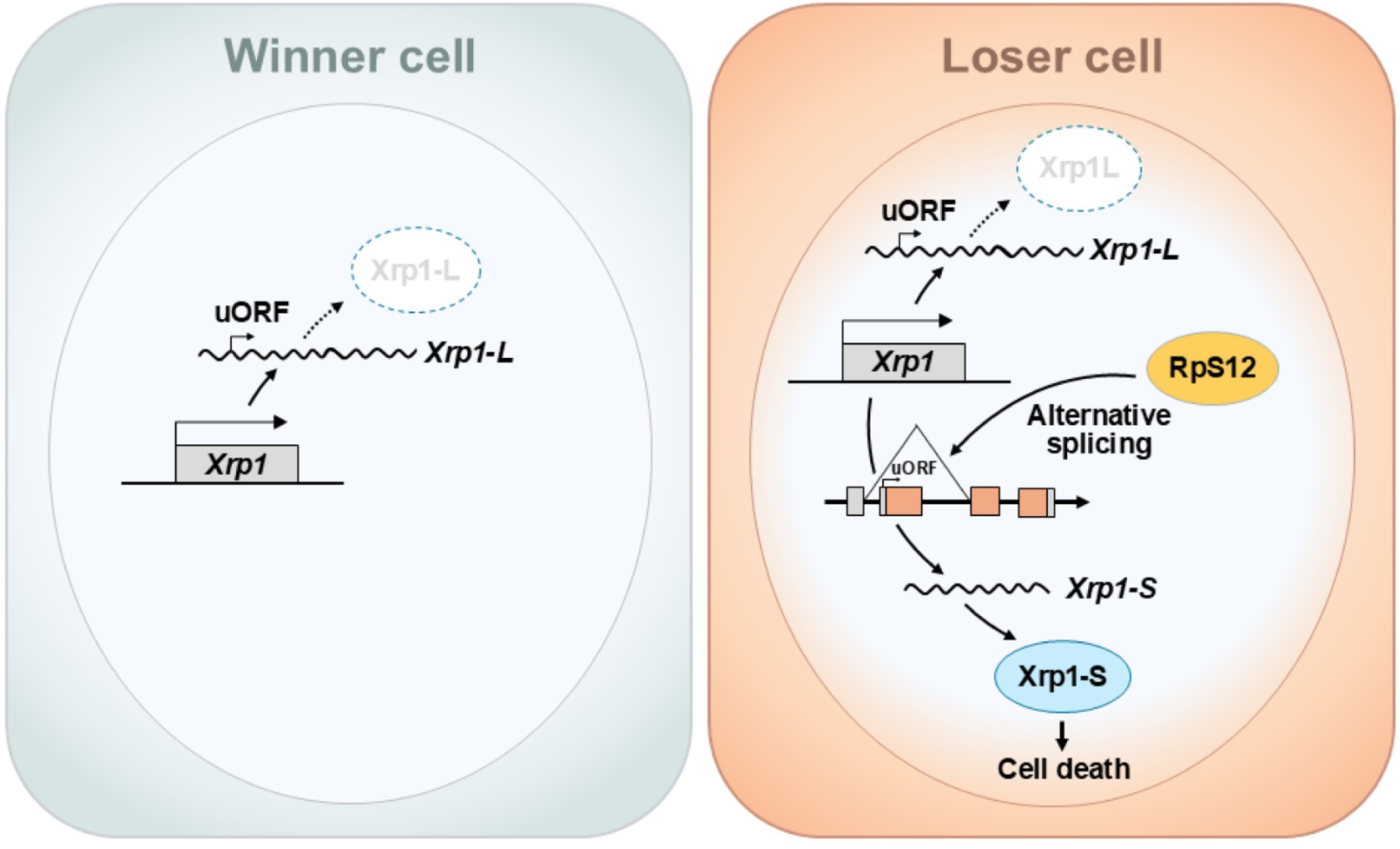
A model for RpS12-mediated Xrp1 induction in loser cells. A schematic representation of Xrp1 induction in unfit cells. RpS12-mediated splicing of *Xrp1* mRNA removes uORF, thereby unleashing Xrp1 translation and causing cell death. See text for details.

To further validate our findings, we disrupted the endogenous Xu288 start codon using the cytosine base editor (CBE) (Huang, Newby and Liu, 2021; Doll, Boutros and Port, 2023), which causes site-specific C to T conversion at the antisense strand and thus G to A conversion at the sense strand of the Xu288 ATG codon, thereby leading to ATG to ATA conversion in S2 cells (Figure 4H, Figure S5) (see Methods). Significantly, disrupting the Xu288 start codon using the guide RNA causing ATG to ATA conversion (gRNA^Xu288-ATA^) resulted in generation of Xrp1-L protein in S2 cells (Figure 4I, Figure S5). This demonstrates that Xrp1 translation is normally repressed by Xu288 translation and thus removal of Xu288 translation unleashes Xrp1 translation to cause cell death.

Finally, we confirmed this notion in the wing disc *in vivo* by inducing phosphorylation of eIF2α, which causes leaky ATG scanning by the ribosome, resulting in skipping the first start codon and thereby increasing translation initiation at the next, downstream start codon (Wek, 2018). Thus, induction of eIF2α phosphorylation would cause skipping of Xu288 start codon and therefore increases translation of downstream Xrp1-L main ORF. Significantly, induction of eIF2α phosphorylation by depleting GADD34, an eIF2α phosphatase, in the wing disc resulted in generation of Xrp1-L protein (Figure 4J). Together, these data indicate that translation of Xrp1-L is normally repressed by a uORF present in the *Xrp1-L* mRNA, whereas unfit cells increase the expression of the *Xrp1-S* variant, which lacks the uORF, resulting in translation of Xrp1-S and subsequent cell death.

## Discussion

Cell competition is initiated upon elevation of intracellular Xrp1 levels in unfit cells, although the underlying mechanism of how Xrp1 is upregulated has long been elusive. In this study, we have uncovered that Xrp1 is abundantly transcribed but is not translated in normal cells due to the presence of uORF; however, in unfit cells, RpS12 causes alternative splicing of *Xrp1-L* mRNA to increase *Xrp1-S* variant, which lacks uORF and is thus translated to Xrp1-S protein that causes cell death. To the best of our knowledge, this post-transcriptional regulation of translation via selective removal of uORF represents a novel way of upregulating protein levels. A similar post-transcriptional upregulation of a gene product can be seen in cells undergoing endoplasmic reticulum (ER) stress, in which a specific, cytoplasmic splicing of *X-box binding protein 1* (*Xbp1*) mRNA results in a production of functional Xbp1 transcription factor (Yoshida *et al*., 2001). Given that the context-dependent splicing of *Xbp1* mRNA is evolutionarily conserved, our finding that ribosomal protein causes upregulation of a stress-responsive protein via uORF removal may potentially be conserved throughout evolution. Notably, the regulation of Xrp1 is strikingly reminiscent of vertebrate CHOP/Ddit3, a C/EBP-family transcription factor that mediates cell death in response to prolonged activation of the integrated stress response (Oyadomari and Mori, 2004). Like Xrp1, CHOP is transcriptionally induced by ATF4 and translationally controlled by a uORF that is bypassed by eIF2α phosphorylation. These parallels further suggest the convergent evolution in the regulatory strategies governing stress-induced cell elimination between flies and vertebrates.

Our data suggest that the Xrp1 gene normally produces Xu288 protein in wild-type cells while it generates Xrp1-S protein in unfit cells by splicing uORF-containing exon 4 out from *Xrp1* mRNA. Disruption of the endogenous Xu288 start codon results in elevated levels of Xrp1-L protein (Figure 4) and increased cell death (data not shown). These observations raise the possibility that normally translated Xu288 has a different function from Xrp1, as was seen in other genes such as PTEN, Myc, and PKC-η genes (Huang *et al*., 2021; Jayaram *et al*., 2021; Li *et al*., 2024). Xrp1 and Xu288 may also serve as complementary *in vivo* markers of cell competition. While both Xrp1-L and Xrp1-S could induce cell death (Figure S3), intriguingly it has also been reported that Xrp1-L affects animal viability more severely than Xrp1-S (Mallik *et al*., 2018). Thus, it would be interesting to investigate the roles of Xrp1-L and Xrp1-S in different types of cell competition. Since Xrp1-L protein is induced by eIF2α phosphorylation, which is provoked by ER stress—a known trigger of cell competition (Ochi *et al*., 2021)—it is possible that distinct triggers of cell competition utilize different Xrp1 isoforms to execute loser’s death. Intriguingly, *Minute* cells undergo proteotoxic stress (Baumgartner *et al*., 2021; Recasens-Alvarez *et al*., 2021)) which was shown to be a driver of cell competition (Baumgartner *et al*., 2021), suggesting that *Minute* cell competition is also driven, at least in part, by eIF2α phosphorylation-mediated induction of Xrp1-L protein. Indeed, this is consistent with our data that *ΔXrp1-L* mutation partially suppressed *Minute* cell competition.

A remaining key question is how RpS12 regulates the alternative splicing of Xrp1 mRNA. Notably, beyond their canonical roles in translation, several ribosomal proteins, such as RpL10A, RpS13, RpS16, and RpS26, regulate their own expression by modulating the splicing of their own mRNAs (Ivanov, Malygin and Karpova, 2005; Malygin *et al*., 2007; Ivanov *et al*., 2010; Takei *et al*., 2016). Moreover, a ribosomal protein RpL22, by cooperating with a splicing regulator hnRNP-A1, regulates alternative splicing of *smad2* mRNA to produce an exon 9-skipped variant, which controls zebrafish morphogenesis (Zhang *et al*., 2017). Given its structural similarity to the spliceosomal component SNU13/NHP2L1, RpS12 may similarly regulate the splicing of *Xrp1* mRNA by cooperating with other splicing regulators.

Our finding that a ribosomal protein induces intracellular accumulation of Xrp1 is reminiscent of the established mechanism by which p53 is upregulated via ribosomal protein-mediated post-transcriptional regulation. In mammals, cellular stresses such as DNA damage and oxidative stress induce nucleolar stress, which leads to the release of free ribosomal proteins (RPs) such as RpL5 and RpL11 from the nucleolus to the nucleoplasm, where they inhibit the E3 ligase MDM2, thereby stabilizing p53. Notably, despite their structural differences, *Drosophila* Xrp1 and mammalian p53 exhibit remarkable functional similarities: both are upregulated in response to DNA damage, regulate overlapping sets of target genes (Khan and Baker, 2022), and play pivotal roles in driving cell competition (Baker, 2020). Future studies investigating the upstream events that enable RpS12 to regulate *Xrp1* mRNA splicing will help elucidate the entire mechanism triggering cell competition—that is, the molecular basis of how cellular fitness is determined.

## Acknowledgments

The authors thank M. Enomoto, D. Kitamura, Y. Oda, D. Morito, S. Yoshimura, S. Sekine and Y. Sekine for discussions; M. Matsuoka, M. Koijima and M. Sada for technical support; S. Goulas for valuable comments on the manuscript; M. Kiparaki for sharing information prior to submission; F. Port and M. Boutros for providing plasmids for the base-editing; E. Piddini, N. Baker, E. Storkebaum, K. Basler, the Bloomington Drosophila Stock Center (BDSC, Indiana, USA), the Vienna Drosophila Resource Center (VDRC, Vienna, Austria), the Drosophila Genomics and Genetic Resources (DGGR, Kyoto, Japan), and National Institute of Genetics (NIG, Shizuoka, Japan) for providing fly stocks. This work was supported in part by grants from the MEXT/JSPS KAKENHI (24K23207 to B.K, 24K22067 to H.K, 21H05284 and 21H05039 to T.I), AMED-CREST, Japan Agency for Medical Research and Development (23gm1710002h0002) to T.I, and the Takeda Science Foundation to H.K and T.I.

## Author contributions

B.K., H.K., R.M., Ki.T. and T.I. designed experiments; B.K., H.K., R.M., S.U., and R.N. conducted experiments with input from T.I.; K.M., K.S. and S.K. generated *Xrp1^040^ and Xrp1^041^* mutant flies; S.Y., Ka.T. and Y.M. conducted RNA-seq; T.K. and M.M. generated Xrp1 antibodies; B.K., H.K., R.M., S.Y., Ka.T. and Y.M. and T.I. analyzed the data; B.K. and T.I. wrote the manuscript; All authors reviewed the manuscript.

## Ethics declarations

The authors declare no competing interests.

## Data availability

All data reported in this paper will be shared by the lead contact upon request. The raw and processed RNA-seq data generated in this study will be deposited in the NCBI Gene Expression Omnibus (GEO); the accession codes will be available before publication. Source data behind the graph is provided in Supplementary Data.

## Code availability

No custom code was used in this study.

## Material & Methods

### Fly strains and generation of clones

*Drosophila melanogaster* strains were raised at 25 °C in vials containing a standard cornmeal-yeast food. For the generation of mitotic clones with gene mutations or transgenes, 20-30 virgin females of healthy tester strains were crossed with approximately 10 healthy males, then transferred to new vials for subsequent egg-laying. Egg-laying was allowed for 8-12 hours to synchronize the developmental stage of the third instar larvae. Fluorescently-labeled mitotic clones were induced into larval eye-antenna imaginal discs or wing imaginal discs. Both males and females at the wandering third instar larval stage were collected for each assay, except in analyses of flies that bore transgenes on the X chromosome. For the generation of clones using inducible hs-FLP, the heat shock induction timing was differentiated depending on the genotype of the tester. For induction of *RpS3^-/+^* cells with *hsFLP RpS3* tester: larvae were subjected to 25 min heat shock at 37 °C, 52 ± 4 hours after egg laying (AEL), then dissected 72 hr later after clone induction. For induction of *RpL19^-/+^* cells with *hsFLP RpL19* tester: larvae were subjected to 25 min heat shock at 37 °C, 60 ± 6 hours after egg laying (AEL), then dissected 72 hr later.

Mosaic clones were induced with strains as follows: *hsFLP; act5C>CD2>GAL4, UAS-mCherry-CAAX, Df(2R)ME; 82B, UbiGFP, TG80, M{RpL19 genomic} (hsFLP RpL19 tester,* a kind gift from N. Baker), *hsFLP,UAS-CD8::GFP;;RpS3[plac92], act>RpS3>GAL4/TM6 (hsFLP RpS3 tester,* a kind gift from E. Piddini), and *w*; tub-Gal80, FRT40A; eyFLP, Act>y+>Gal4, UAS-GFP (40A MARCM tester*). Other strains used for analyses are as follows: *Xrp1^m2-73^* (a kind gift from N. Baker), *RpS12^G97D^* (a kind gift from N. Baker), *UAS-RpS12* (a kind gift from N. Baker), *RpS18M(2)56-f* (a kind gift from N. Baker)*, Xrp1^08^* (a kind gift from K. Basler), *UAS-Xrp1^Long^* (a kind gift from E. Storkebaum), *UAS-Xrp1^Short^* (a kind gift from E. Storkebaum), *Xrp1^040^ (Xrp1^ΔLong^),* (This study, see Methods), *Xrp1^041^ (Xrp1^ΔLong^)* (This study, see Methods), *UAS-Xrp1-RNAi^TRiP.HMS00053^* (BDSC, #34521), *UAS-PPP1R15-RNAi^HMS00811^* (BDSC #33011), *UAS-ATF4* (BDSC, #81655), *Xrp1 ^02515^ (Xrp1-lacZ) (BDSC, #11569), Hel25E^ccp-8^* (Nagata *et al*., 2019), *RpS3^2^* (BDSC, #5627), *UAS-Luc-RNAi^TRiP.JF01355^* (BDSC#31603), *UAS-Luc* (BDSC, #35788). Detailed genotypes used in this study are provided in Supplementary Table 1.

### Immunohistochemistry

Wandering third-instar larvae were dissected in PBS, and fixed in 4 % paraformaldehyde for 20 min at RT. Then samples were washed three times in PBT (PBS + 0.1 % Triton X-100). Tissues were then blocked in PBT containing 5 % normal donkey serum (Jackson ImmunoResearch, #107-000-121) and incubated at 4 °C with primary antibodies.

Primary antibodies used are as follows: rat anti-Xrp1 (This study, see “Generation of Xrp1 monoclonal antibody” in Methods section, 1:50), chicken anti-β-galactosidase (Abcam, # ab9361, 1:1000), rabbit anti-cleaved Drosophila Dcp1 (Asp216) (Cell Signaling Technology, # 9578, 1:100), Secondary antibodies used are as follows: Goat anti-Rabbit IgG (H+L) Highly Cross-Adsorbed Secondary Antibody, Alexa Fluor™ 546 (Thermo Fisher Scientific, #A11035, 1:250), Goat anti-Rat IgG (H+L) Highly Cross-Adsorbed Secondary Antibody, Alexa Fluor™ Plus 555 (Thermo Fisher Scientific, # A48263, 1:250), Goat anti-Chicken IgY (H+L) Cross-Adsorbed Secondary Antibody, Alexa Fluor™ Plus 555 (Thermo Fisher Scientific, # A32932, 1:250), Goat anti-Rat IgG (H+L) Highly Cross-Adsorbed Secondary Antibody, Alexa Fluor™ Plus 647 (Thermo Fisher Scientific, # A48265, 1:250).

### Generation of *Xrp1^040^* and *Xrp1^041^* mutant flies

The mutant alleles *Xrp1^040^ and Xrp1^041^* were generated using the CRISPR/Cas9 system, as previously described (Kondo and Ueda, 2013). The guide RNA sequence used was 5ʹ-AACTCGTTGCAGGCGCGGCC-3ʹ.

### Generation of Xrp1 monoclonal antibody

His-tagged (for antigen production) and MBP-tagged (for monoclonal antibody screening) full-length Xrp1 were expressed in BL21-CodonPlus-RP (Agilent Technologies, Santa Clara, CA) transformed with pET-28a (Invitrogen) and pMAL (New England Biolabs, Beverly, MA), respectively. Fusion proteins were purified by affinity chromatography on TALON metal-affinity resin (Clontech) for His tags or amylose resin (New England Biolabs) for MBP tags. A rat monoclonal antibody against Xrp1 was generated as described previously (Kobayashi *et al*., 2022). Briefly, WKY/NCrl rats were immunised with an emulsion of purified Xrp1 antigen. Twenty-one days after the injection, splenic lymphocytes were fused with SP2/0-Ag14 myeloma cells. Hybridoma culture supernatants were screened for Xrp1 reactivity by solid-phase ELISA, and positive clones were selected for expansion.

### Cloning and plasmid construction

Amplicons were obtained by PCR reaction using KOD One® PCR Master Mix (TOYOBO, # KMM-101) and primers containing the corresponding sequence for In-Fusion reaction. 5’UTR of Xrp1-RA were amplified cDNA extracted from *w^1118^*flies. CDS of mScarlet-I was amplified from pF3BGX-mScarlet-I (addgene, #138391). CDS of 2 isoforms of Xrp1 was respectively amplified from pBSSK-Xrp1 (Long isoform)-PstI, which was used for the generation of Xrp1 antibody. To construct Xrp1-RA_5’UTRuATGdel-mScarlet-I, amplicons were amplified from the pUASTattB-Xrp1-RA_5’UTR-mScarlet-I plasmid. Each amplicon was cloned into XbaI-digested pUAST-attB (Drosophila Genetic Resource Center, #1419) using the In-Fusion HD Cloning Kit (Clontech, #639648). The primer sequences used for PCR are as follows:

Xrp1^long^-fw: 5′-CGAGGGTACCTCTAGCAAAAATGATCCAGGAGCCAGCAC –3′

Xrp1^long^-rv: 5′-ACAAAGATCCTCTAGTCAGTCCTGCTCCTGCTTAACGT –3′

Xrp1^short^-fw: 5′-CGAGGGTACCTCTAGCAAAAATGTTTGCCGAGGAGGATCT –3′

Xrp1^short-^-rv: 5′-ACAAAGATCCTCTAGTCAGTCCTGCTCCTGCTTAACGT –3′

Xrp1-RA_5’UTR-fw: 5′-CGAGGGTACCTCTAGTCATTTCTGTTCGCCACTA –3′

Xrp1-RA_5’UTR-rv: 5′-CAATATTATATCTCTGGGATTTTCGGT –3′

mScarlet-I-fw: 5′-AGAGATATAATATTGATGGTGAGCAAGGGCGAGGCAGTGAT –3′

mScarlet-I-rv: 5′-ACAAAGATCCTCTAGTTACTTGTACAGCTCGTCCAT –3′

Xrp1-RA_5’UTR^uATGdel^ fragment1-fw: 5′-CGAGGGTACCTCTAGTCATTTCTGTTCGCCACTA –3′

Xrp1-RA_5’UTR^uATGdel^ fragment1-rv: 5′-TTCGACGGAGAATGCTTTGAACGTGGTTTTAATTG –3′

Xrp1-RA_5’UTR^uATGdel^-mScarlet-I fragment2-fw: 5′-GCATTCTCCGTCGAAGCCCAAACCA –3′

Xrp1-RA_5’UTR^uATGdel^-mScarlet-I fragment2-rv: 5′-ACAAAGATCCTCTAGTTACTTGTACAGCTCGTCCAT –3′

### Base editing with CBE^evoCDA1^

gRNA target site was predicted by using CRISPOR (https://crispor.gi.ucsc.edu/) against the *D. melanogaster* dm6 reference genome. The pCFD5 plasmid (Port and Bullock, 2016) (a kind gift from F. Port) was linearized with BbsI, and annealed oligos were inserted using the In-Fusion HD Cloning Kit (Clontech, #639648). For base editing, S2 cells were co-transfected with pAct-CBE^evoCDA1^ (Doll, Boutros and Port, 2023) (a kind gift from F. Port) and the corresponding pCFD5-gRNA plasmid. Genomic DNA was extracted from S2 cells with the QIAamp DNA Micro Kit (QIAGEN, #56304). A genomic fragment encompassing the Xu288 region was then amplified by PCR with the primers listed below. Editing efficiency was calculated by using the base editing analysis tool (BEAT) (Xu, Liu and Han, 2019)(https://hanlab.cc/beat/). Oligos used are as follows:

gRNA^Xu288ATA^-fw: 5′-TTCCCGGCCGATGCAGGAGAATGCCATTTTGAACG-3′

gRNA^Xu288ATA^-rv: 5′-TTCTAGCTCTAAAACCGTTCAAAATGGCATTCTCC-3′

Primers used for the amplification of Xu288 region and sequencing are as follows:

Xu288-fw: 5′-GATATCCTGTCGATCCGAAACCG-3′

Xu288-rv: 5′-GAGAGAGTGCTGGGTACACAGGC-3′

Sequencing-fw: 5′-GATATCCTGTCGATCCGAAACCG-3′

Sequencing-rv: 5′-TGGTTGATTGGGTTGCTGTTGTTTG-3′

### Cell culture and transfection

S2 cells were cultured at 26 °C in Schneider’s *Drosophila* Medium (Gibco, #21720024) supplemented with 10 % heat-inactivated fetal bovine serum (FBS; Sigma, #S173012) and 0.5 % Penicillin–Streptomycin (Sigma, #P4333). Cells were transfected with plasmid DNA using Effectene Transfection Reagent (QIAGEN, #301425).

### Foldseek analysis

Structural homologues of *Drosophila* RpS12 were identified with Foldseek (https://search.foldseek.com). The predicted RpS12 structure (AF-P80455-F1-v4) was downloaded from the AlphaFold Protein Structure Database (https://alphafold.ebi.ac.uk/). Searches were run against the AlphaFold/UniProt database in 3Di/AA mode with the taxonomic filter set to “Homo sapiens”.

### Western Blotting

25 third-instar larvae were dissected in 1× PBS, and 50 wing imaginal discs were collected. Discs were homogenised in 25 µL RIPA buffer (FUJIFILM Wako, #188-02453) plus 25 µL Sample Buffer Solution (Nacalai Tesque, #30566-22) using BioMasher II (Nippi, #893061), then boiled at 95 °C for 5 min. For S2 cells, pellets were resuspended in 200 µL Sample Buffer Solution and boiled at 96 °C for 10 min. Per lane, 20 µL lysate and 2 µL BLUE Star prestained protein ladder (Nippon Genetics, #NE-MWP03) were loaded onto 8 % SDS–PAGE gels. Proteins were transferred to PVDF membranes (Millipore, IPVH00010), blocked with 5% skim milk, and then washed in TBST (TBS + 0.1 % Tween-20). Membranes were incubated overnight at 4 °C with rat anti-Xrp1 (1:100), then probed for 2 h at RT with HRP-conjugated anti-rat IgG (Cell Signaling Technology, #7077; 1:1000). Signals were developed with Chemi-Lumi One Super (Nacalai Tesque, #02230) and imaged on a FUSION SOLO.7S.EDGE system (Vilber). Membranes were stripped with WB Stripping Solution (Nacalai Tesque, #05364-55) and reprobed with mouse anti-α-tubulin (Sigma, #T5168; 1:5000) followed by HRP-conjugated anti-mouse IgG (Cell Signaling Technology, #7076; 1:1000) as a loading control.

### RNA extraction and semi-quantitative RT-PCR

Wing imaginal discs were dissected from wandering third-instar larvae, and total RNA was extracted with the RNeasy Mini Kit (QIAGEN, #74104) according to the manufacturer’s instructions. First-strand cDNA was synthesized from the purified RNA with the SuperScript IV First-Strand Synthesis System (Invitrogen, #18091050). PCR reaction was performed with GoTaq Green Master Mix (Promega, #M7123). PCR conditions were as follows: 95°C for 2min, 38 cycles of (95°C for 1 min, 50°C for 45 sec, 72°C for 1 min) and 72°C for 1 min. Primers used are as follows:

Primer-set 1 (for amplification of Xrp1-RA, RB, RC, RG)

semiv1-fw: 5′-GAACTTGTGTCGACGACTCTTTTAG –3′

semiv1-rv: 5′-GCGAGTTGTGGAATATGTTGATGAA –3′

Primer-set 2 (for amplification of Xrp1-RA, RC, RE, RG)

semiv2-fw: 5′-CGGTTGACTCAGAGATCGCA –3′

semiv2-rv: 5′-GCTCCTGGCTGCTGGTATAG –3′

Primer-set 3 (for amplification of Xrp1-RD)

semiv3-fw: 5′-AAGAGGTGTTTTTGGTCCGC –3′

semiv3-rv: 5′-GTGTGGTCGGACTGAATGCT –3′

### RNA-seq

Detailed methods for CAGE library preparation, generation of CAGE tag clusters, preparation of Oxford Nanopore Technologies long-read RNA-seq library, sequencing, and processing of long-read RNA-seq data will be published elsewhere.

### Image acquisition

Confocal images were acquired with a confocal microscope ZEISS LSM 880 (Carl Zeiss) under the control of ZEN Blue (Carl Zeiss) or Laser Scanning Confocal Microscope TCS SP8 on DMi8 inverted microscope (Leica Microsystems) under the control of Leica Application Suite X version 3.5.5.19976 (Leica Microsystems) or STELLARIS 5 WLL on DMi8 inverted microscope (Leica Microsystems).

### Quantification

XY confocal images of wing imaginal discs stained with the indicated antibodies and DAPI, as well as agarose gel images from semi-quantitative RT-PCR, were processed in FIJI (ImageJ-win64 v1.54f). Subsequent analysis was differentiated depending on the antibodies used for staining.

### Statistics and Reproducibility

All statistical analyses were carried out in R (v4.4.1) within RStudio (v2024.12.1). Significances are shown as follows. *: p<0.05, **: p<0.01 ***: p<0.001, n.s. (not significant): p>0.05. For single comparisons, Wilcoxon rank sum test or Welch’s t-test was performed. For multiple comparisons, one-way ANOVA followed by Tukey’s HSD test or Kruskal–Wallis test followed by the Steel–Dwass test were performed, as appropriate. Details of statistical evaluations and the sample sizes were written in the Figure legends. Sample size adjustment was not performed by predetermining test. Each data point in the plots represents an independent biological replicate, and each experiment was performed independently at least three times.

## Figure Legends

**Supplementary Figure 1.**
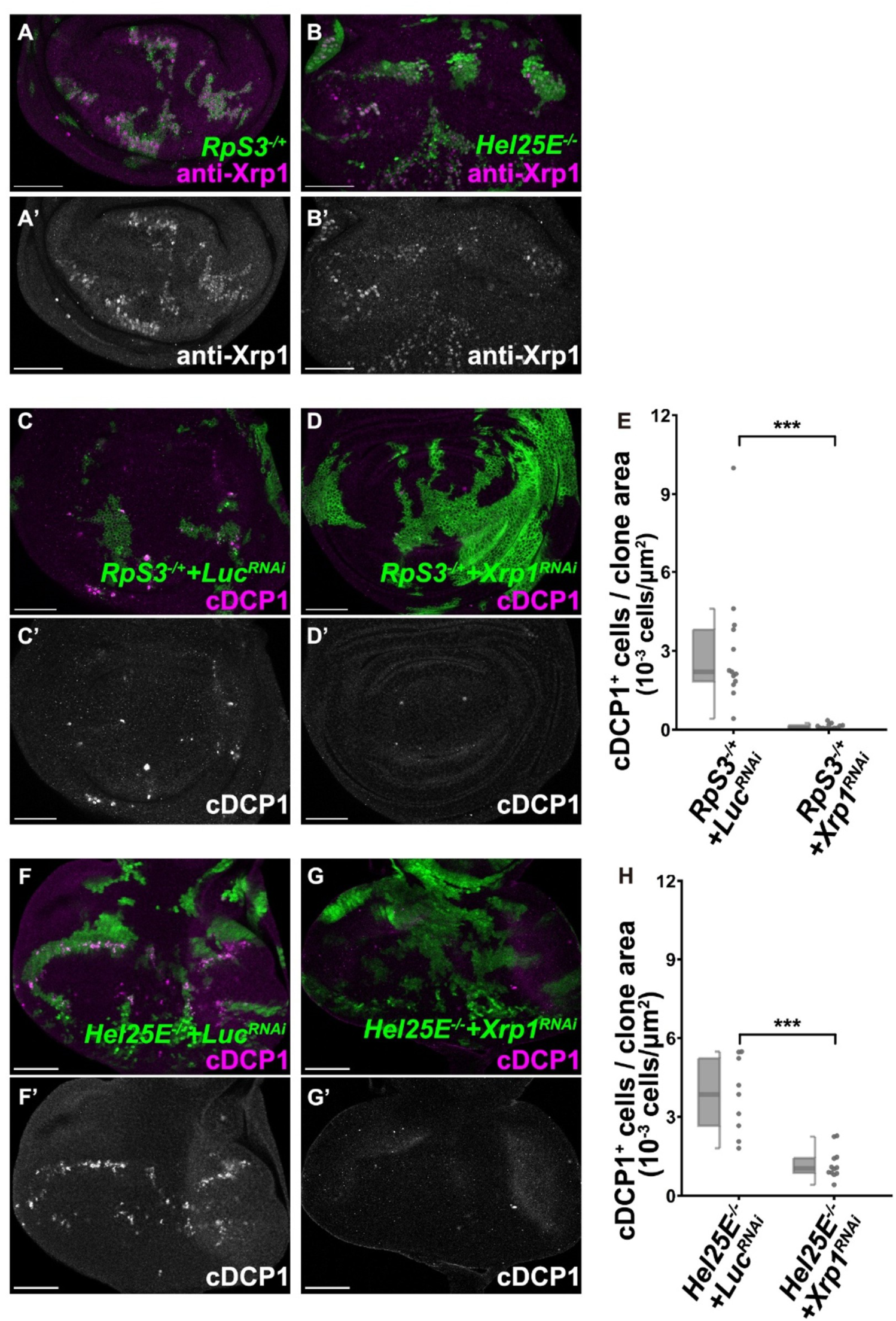
Xrp1 induction is required for cell competition. **(A)** Wing disc bearing *hsFLP* induced *RpS3^-/+^*cells stained with anti-Xrp1 antibody.. Scale bars, 50μm. (B) Eye disc bearing *eyFLP* induced *Hel25E*^-/-^ clones stained with anti-Xrp1 antibody. Scale bars, 50μm. (C,D) Wing disc bearing *hsFLP* induced *RpS3^-/+^ +Luc-RNAi* (C) or *RpS3^-/+^ +Xrp1-RNAi* (D)cells stained with anti-cDCP1 antibody. Scale bars, 50μm. (E) Boxplot overlaid with dot plot represents the number of c-Dcp1 positive cells in GFP-positive region for *RpS3^-/+^ +Luc-RNAi* (n=13), *RpS3^-/+^ +Xrp1-RNAi* (n=13) samples. Each plot corresponds to the raw data. Statistical significance is shown as follows: ***p < 0.001; Wilcoxon rank sum test. (F,G) Eye disc bearing *eyFLP* induced *Hel25E^-/-^ +Luc-RNAi* (F) or *Hel25E^-/-^ +Xrp1-RNAi* (G)cells stained with anti-cDCP1 antibody. Scale bars, 50 μm. (H) Boxplot overlaid with dot plot represents the number of c-Dcp1 positive cells in GFP-positive region for *Hel25E^-/-^ +Luc-RNAi* (n=9) or *Hel25E^-/-^ +Xrp1-RNAi* (n=12) samples. Each plot corresponds to the raw data. Statistical significance is shown as follows: ***p < 0.001; Wilcoxon rank sum test.

**Supplementary Figure 2.**
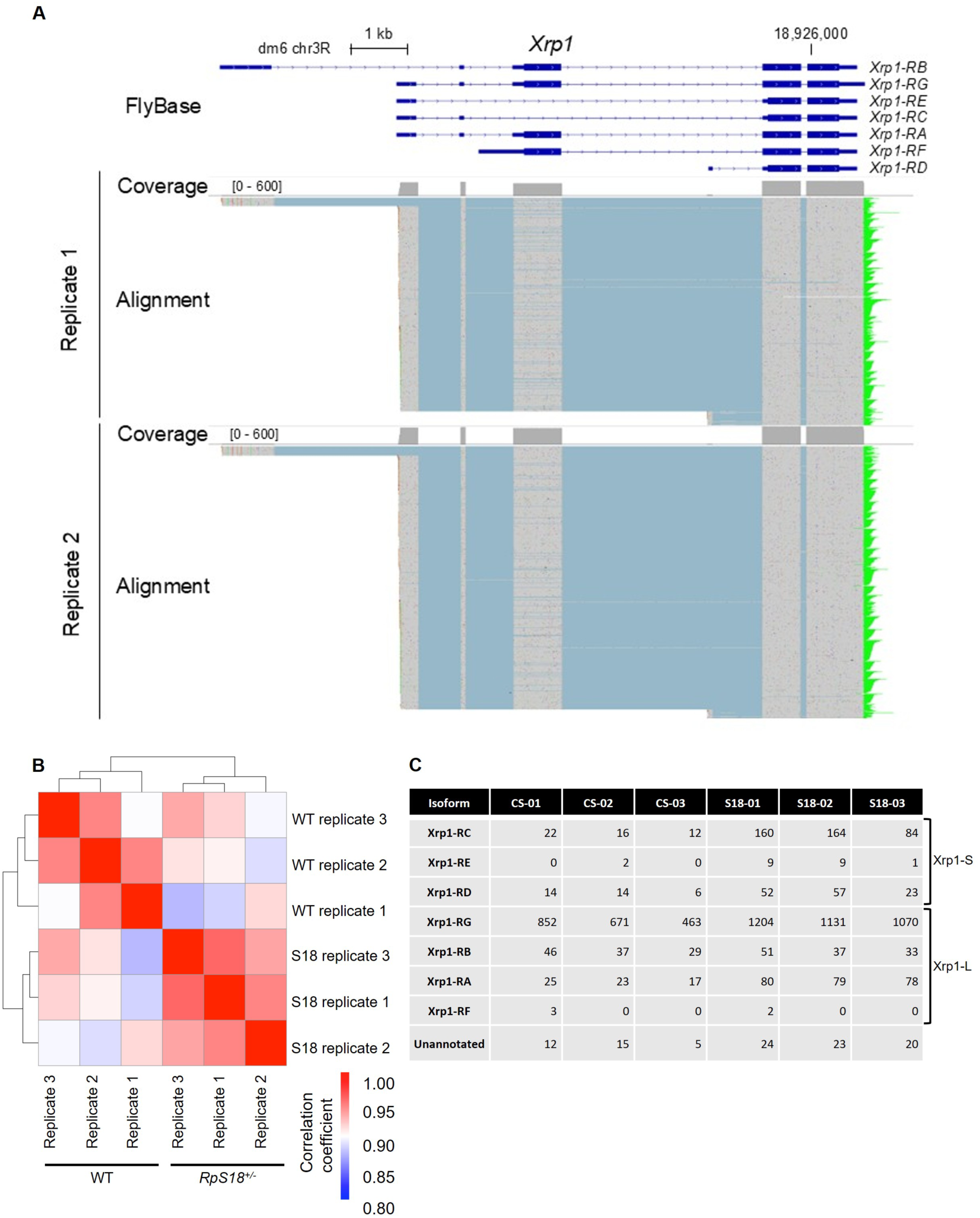
*Xrp1* mRNA isoforms revealed by full-length RNA-seq. (A) Genome browser view of full-length RNA-seq data at the *Xrp-1* locus. *Top*: Seven annotated isoforms of *Xrp1* (*Xrp1-RA, Xrp1-RB, Xrp1-RC, Xrp1-RD, Xrp1-RE, Xrp1-RF and Xrp1-RG*) from the FlyBase dataset (dmel-all-r6.58.gtf). Bottom, IGV snapshot of long-read RNA-seq data at the *Xrp-1* locus in two control replicates, shown on the forward strand. (B) A heatmap of Pearson correlation coefficients was generated from log2-counts per million (CPM) values for genes based on the FlyBase dmel-all-r6.58.gtf dataset. The analysis reveals high correlation among biological replicates within the wild-type (WT) and *RpS18^+/-^* groups, indicating high reproducibility of the experiment. (C) Raw counts of Xrp1-isoforms in wing discs from wild-type (WT) and *RpS18^+/-^*.

**Supplementary Figure 3.**
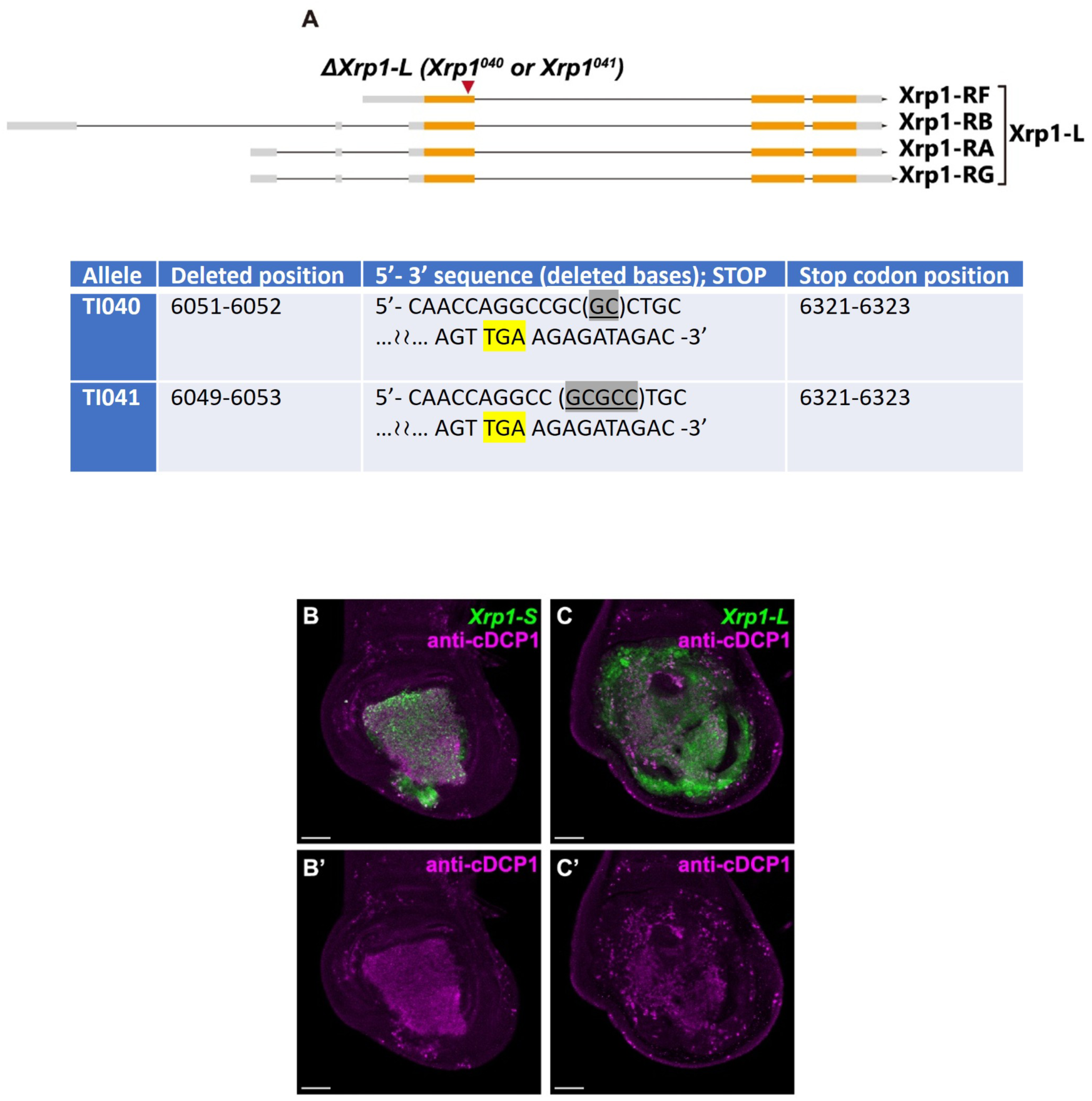
Mutations and functions of Xrp1-L and Xrp1-S. (A) The detailed information of *Xrp1^040^* and *Xrp1^041^* alleles. Deleted sequences are highlighted in gray, and the STOP codon is highlighted in yellow. (B, C) Wing discs expressing Xrp1-S (B) or Xrp1-L (C) under *nub-Gal4*, stained with anti-cDCP1 antibody. Scale bars, 50 μm. Confocal images were acquired with Laser Scanning Confocal Microscope STELLARIS 5 WLL on DMi8 inverted microscope (Leica Microsystems).

**Supplementary Figure 4.**
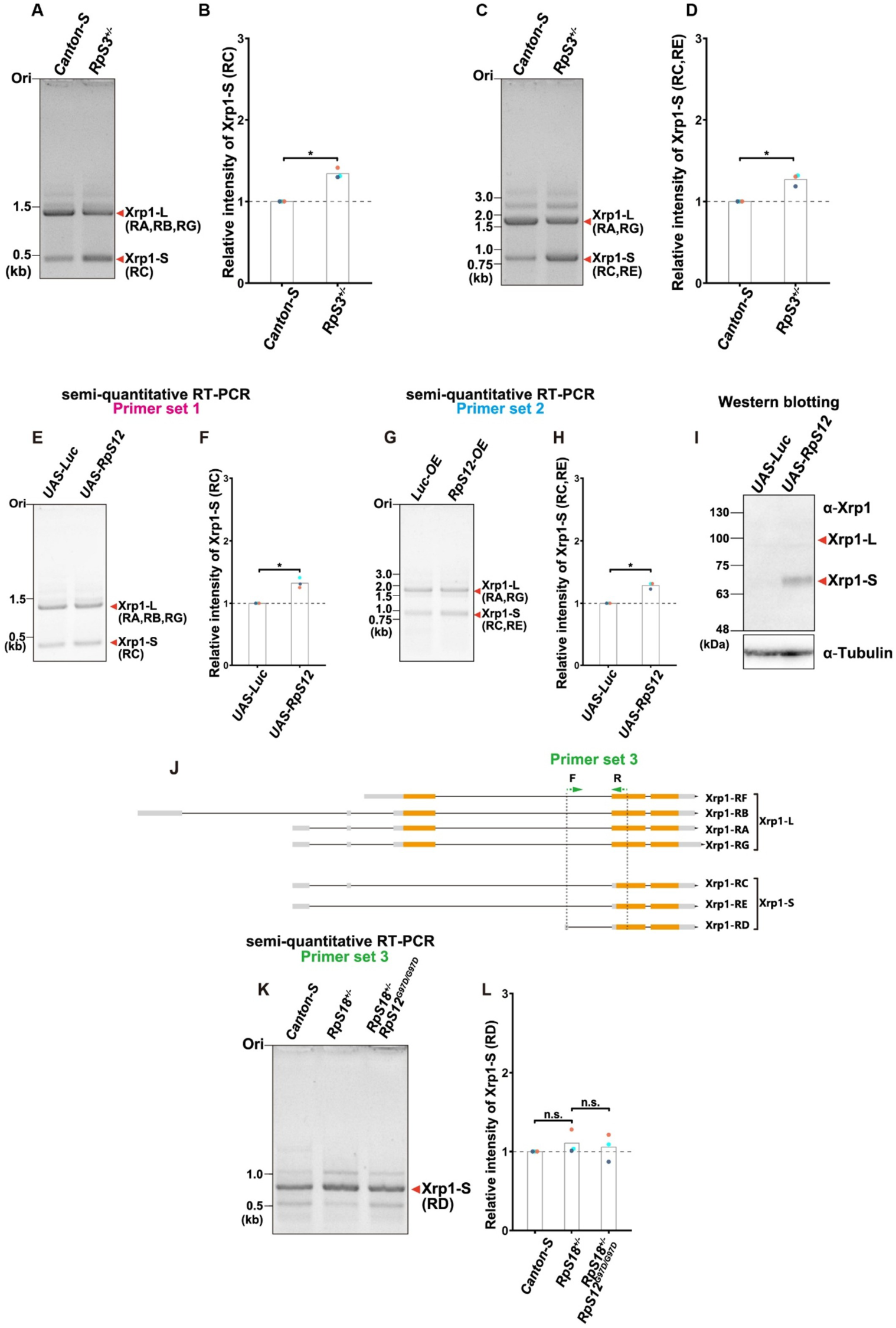
Semi-quantitative PCR for *Xrp1-L* and *Xrp1-S*. (A) Semi-qRT-PCR products amplified using primer set 1 from cDNA derived from *Canton-S* (left) and *RpS3^-/+^*(right). (B) Bar plot overlaid with dot plot represents the relative intensity of *Xrp1-S* (RC) normalized by control sample. Each dot plot corresponds to the raw data, and each plot color corresponds to each replicate. White bars show the average. Statistical significance is shown as follows: *p < 0.05; Welch’s t-test. (C) Semi-qRT-PCR products amplified using primer set 2 from cDNA derived from *Canton-S* (left) and *RpS3^-/+^*(right). (D) Bar plot overlaid with dot plot represents the relative intensity of *Xrp1-S* (RC,RE) normalized by control sample. Each dot plot corresponds to the raw data, and each plot color corresponds to each replicate. White bars show the average. Statistical significance is shown as follows: *p < 0.05; Welch’s t-test. (E) Semi-qRT-PCR products amplified using primer set 1 from cDNA derived from *Luc-OE* sample (left) and *RpS12-OE* sample (right). (F) Bar plot overlaid with dot plot represents the relative intensity of *Xrp1-S* (RC) normalized by control sample. Each dot plot corresponds to the raw data, and each plot color corresponds to each replicate. White bars show the average. Statistical significance is shown as follows: *p < 0.05; Welch’s t-test. (G) Semi-qRT-PCR products amplified using primer set 2 from cDNA derived from *Luc-OE* sample (left) and *RpS12-OE* sample (right). (H) Bar plot overlaid with dot plot represents the relative intensity of *Xrp1-S* (RC, RE) normalized by control sample. Each dot plot corresponds to the raw data, and each plot color corresponds to each replicate. White bars show the average. Statistical significance is shown as follows: *p < 0.05; Welch’s t-test. (I) Wing discs of third instar larvae of each genotype were subjected to Western blot analysis using anti-Xrp1 antibody or anti-Tubulin antibody. (J) Illustration for primer-sets used for the amplification of Xrp1 mRNA. (K) semi-qRT-PCR products amplified using primer set 3 from cDNA derived from *Canton-S* (left), *RpS18^+/-^* (center), and *RpS18^+/-^* with *RpS12^G97D/G97D^*(right). (L) Bar plot overlaid with dot plot represents the relative intensity of *Xrp1-S* (RD) normalized by control sample. Each dot plot corresponds to the raw data, and each plot color corresponds to each replicate. White bars show the average. Statistical significance is shown as follows: n.s. > 0.05; one-way ANOVA followed by Tukey HSD test.

**Supplementary Figure 5.**
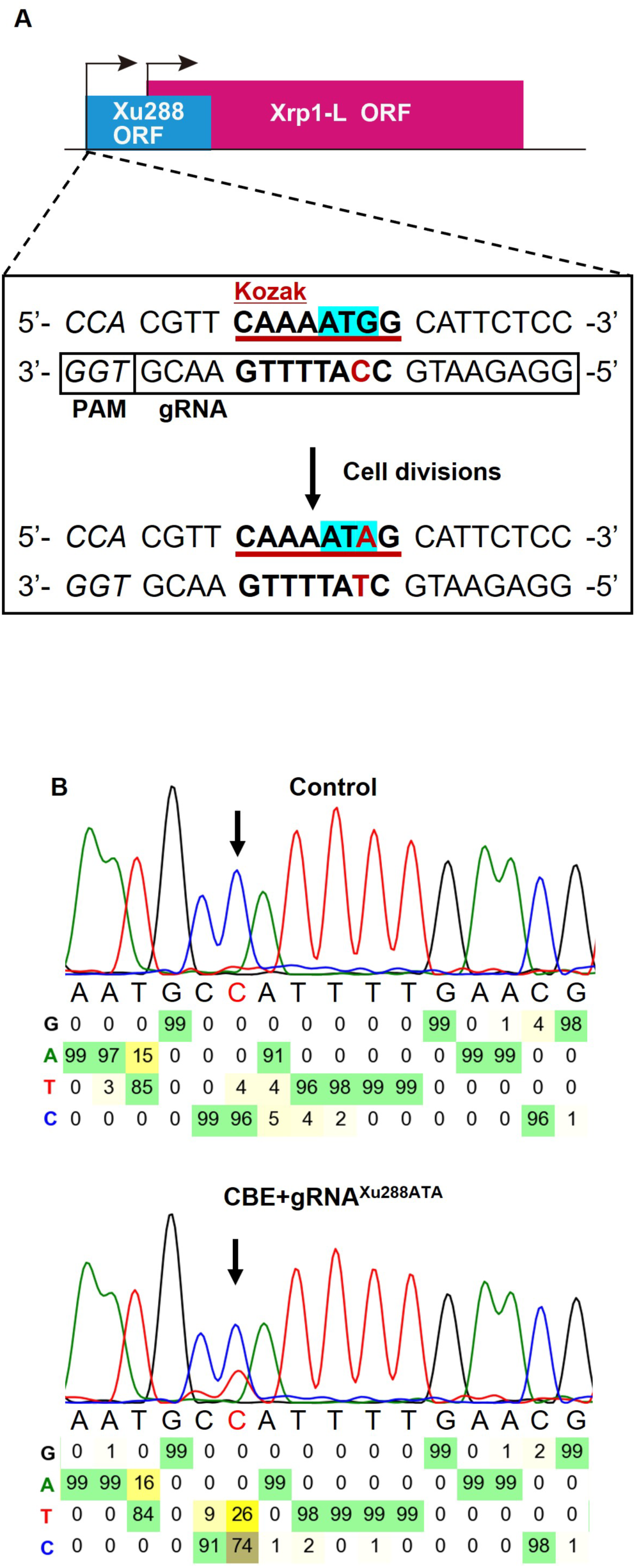
uORF inhibition by cytosine base editor. (A) Schematic of the gRNA^Xu288-ATA^ target site. Kozak motif is underlined.; the original ATG codon of Xu288 is highlighted in blue, and the intended ATA substitution is highlighted in red. (B) Example Sanger sequencing data of genomic DNA from S2 cells co-transfected with CBEevoCDA1 and gRNAXu288-ATA, confirming the ATG→ATA edit.

**Supplementary Table 1:**
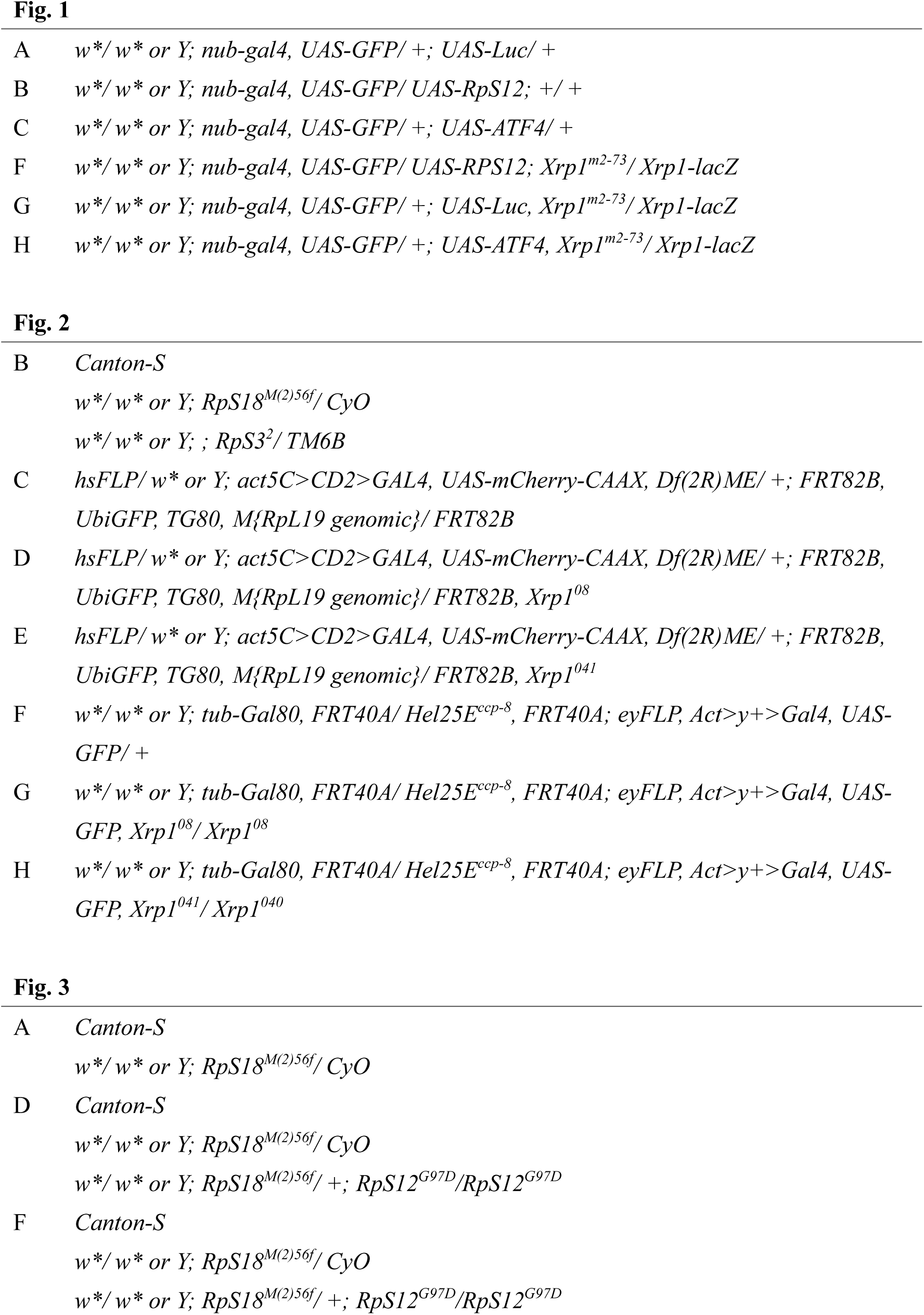
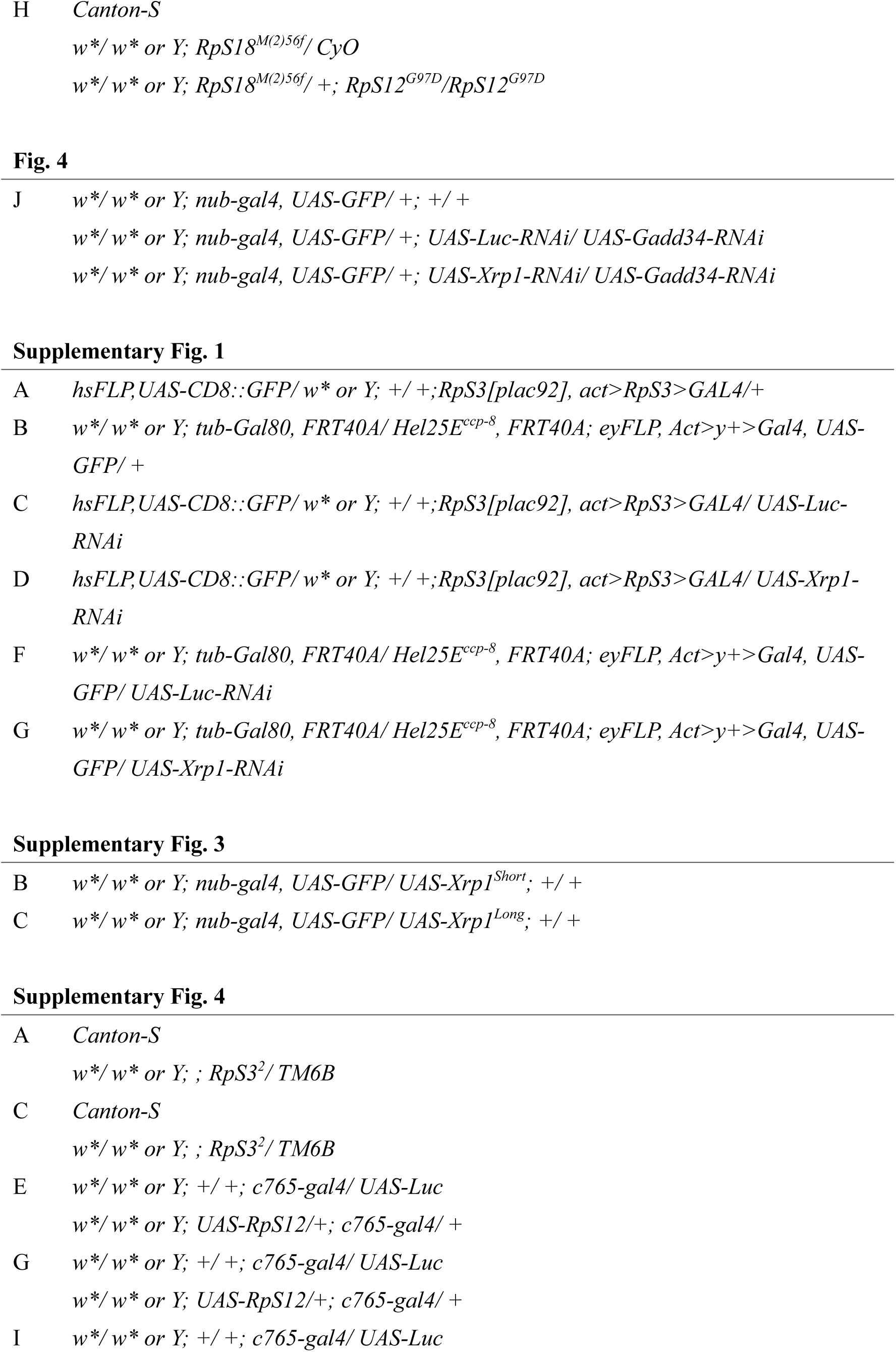
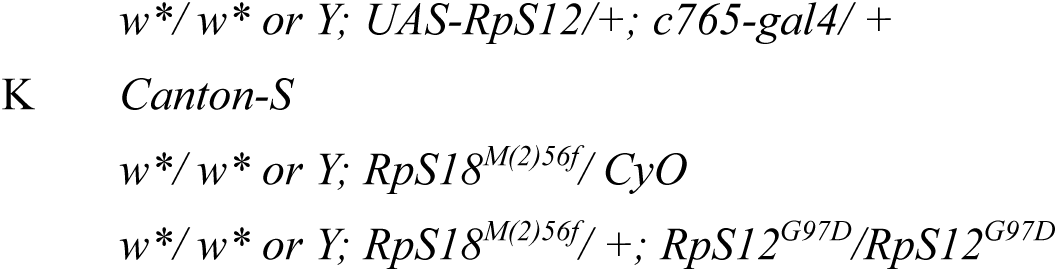
List of genotypes.

